# Male-specific genome sequences for non-invasive sex identification in Nishikigoi (*Cyprinus carpio*)

**DOI:** 10.64898/2026.11.09.765473

**Authors:** Kenta Shirasawa, Masaaki Miyamoto, Kazuto Takasaki

## Abstract

Nishikigoi, ornamental cultivars of the common carp (*Cyprinus carpio*), are economically valuable aquaculture teleosts in which mature females command significantly higher market prices due to superior body conformation and coloration. Consequently, early and non-invasive sex identification is highly desirable for commercial breeding. Although sex determination in common carp follows an XX/XY system, sex-chromosome-specific markers and field-applicable diagnostic methods for Nishikigoi have remained limited. In this study, we conducted sequence-based bulked segregant analysis using whole-genome shotgun sequencing of female and male pools in Kohaku, a representative Nishikigoi lineage. We identified male-specific candidate contigs and successfully developed PCR-based male-specific markers. Validation across multiple Nishikigoi varieties and wild common carp demonstrated that primary primer pairs achieved high diagnostic accuracy. Comparative genomic analysis revealed that these candidate contigs mapped to the genomic region on chromosome 25B containing the male-determining gene *Pangu*, with the primary marker located in immediate physical proximity to the gene. To enable field diagnostics, we combined non-invasive epidermal mucus DNA extraction, loop-mediated isothermal amplification (LAMP), and chromatography-printed array strip (C-PAS) technology, establishing a rapid on- site detection workflow. Overall, this study provides highly reliable PCR markers associated with the sex-determination locus and establishes a non-invasive, field-applicable diagnostic platform to streamline commercial Nishikigoi breeding and sex-selected stock management.

## Introduction

Nishikigoi are ornamental varieties of the common carp (*Cyprinus carpio*). Over two centuries of spontaneous mutation and meticulous selective breeding have established over 100 distinct varieties. These are broadly categorized into major lineages such as the iconic Gosanke (the ‘Big Three’ varieties) comprising Kohaku (white body with red markings), Taisho Sanshoku (white body with red and black markings), and Showa Sanshoku (black body with red and white markings) alongside single-colored or metallic lineages such as Ogon. Among these diverse varieties, female individuals generally command substantially higher market prices and aesthetic value than males, as females develop larger, fuller body conformations and maintain vibrant skin pigmentation at maturity. Consequently, early sex identification is a critical task in commercial breeding. Practical methods for sex identification rely on anatomical and morphological observations of mature individuals, creating a strong demand for non-invasive and reliable diagnostic techniques applicable during juvenile stages and non-breeding seasons.

The common carp exhibits an XX/XY sex-determination system, in which females are homogametic (XX) and males are heterogametic (XY). In common carp, on one hand, genome sequences for both male and female were reported (Chen *et al*. 2026, Xu *et al*. 2014, Yuan *et al*. 2025) and a candidate gene for the sex determination was also identified (Chen *et al*. 2026). On the other hand, chromosomal translocations unique to Nishikigoi compared to wild common carp have been reported (Ojima and Takai 1981), the detailed structure of their sex chromosomes remains poorly understood. Identifying male-specific sequences in Nishikigoi is therefore essential to establish reliable molecular markers for sex identification. However, though the genome sequences for Nishikigoi are publicly available (Hosaka 2023), the genome information was derived from female individuals. Therefore, the sex chromosomes, particularly the Y chromosome, in Nishikigoi are not yet fully characterized.

Bulked segregant analysis (BSA) is a powerful genetic technique used to identify loci associated with specific phenotypic variations (Michelmore *et al*. 1991). When comparing a female bulk pool with a male bulk pool, sex-determining loci are expected to be homozygous (XX) in the female bulk and heterozygous (XY) in the male bulk. Conversely, regions unlinked to sex determination should reflect the background genetic diversity of the population in both bulk pools. With recent advancements in high-throughput DNA sequencing, integrating sequence analysis into BSA (e.g., MutMap or QTL-seq approaches) has enabled rapid and efficient identification of DNA markers tightly linked to target traits (Abe *et al*. 2012, Takagi *et al*. 2013).

Ideally, sex identification in Nishikigoi should be feasible directly in field settings, such as breeding farms and markets, where specialized molecular biology laboratory equipment is unavailable. To meet such on-site diagnostic demands, marker detection systems combining loop-mediated isothermal amplification (LAMP) (Notomi *et al*. 2000) coupled with chromatography-printed array strip (C-PAS) technology (Tohoku Bio Array Co.,Ltd., Sendai, Japan) have been effectively utilized for crop cultivar identification (Monden *et al*. 2014, 2023, Okamoto *et al*. 2023, Takabatake *et al*. 2025) and genetically modified organism detection (Takabatake *et al*. 2018b, 2018a, 2023). Moreover, non-invasive methods are required without harming the fish (Yasuike *et al*. 2026). In this study, we conducted sequence-based BSA to identify male-specific sequences in Nishikigoi. Using these identified sequences, we aimed to develop highly accurate PCR-based sex-identification markers and subsequently establish a non-invasive, field-applicable DNA diagnostic method combining LAMP and C-PAS for sex identification in Nishikigoi.

## Materials and methods

### Ethics Statement

Under the institutional regulations of the Kazusa DNA Research Institute, formal ethical review was not required because fish are not subject to approval by the Animal Experimentation Committee. Nevertheless, all fish were handled in accordance with standard animal care guidelines, with care taken to minimize stress and distress throughout the study.

### Fish samples and sex identification

A total of 240 Nishikigoi individuals, comprising 144 Kohaku, 32 Doitsu Karasu, 32 Aka Bekko, and 32 Ogon, and 60 wild common carp (Magoi) individuals, were used in this study. Sex was identified for each individual by anatomical examination of the gonads following dissection. In addition, we used eight living Ogon individuals for DNA diagnostics.

### DNA extraction and sequencing analysis

Genomic DNA was extracted from approximately 30 mg of pectoral fin tissue from each individual using the DNeasy Blood & Tissue Kit (Qiagen, Hilden, Germany) according to the manufacturer’s instructions. The concentration and purity of the extracted DNA were evaluated using a Qubit Fluorometer with the dsDNA Broad Range Assay Kit (Thermo Fisher Scientific, Waltham, MA, USA) and a NanoDrop

Spectrophotometer (Thermo Fisher Scientific). DNA integrity was assessed via capillary electrophoresis on a TapeStation system (Agilent Technologies, Santa Clara, CA, USA).

To identify male-specific candidate sequences, bulked segregant analysis (BSA) was performed. Genomic DNA from 54 female and 67 male Kohaku individuals were pooled in equimolar amounts to construct a female bulk pool and a male bulk pool, respectively. Whole-genome shotgun libraries were prepared using the Swift 2S TURBO Flexible DNA Library Kit (Swift Biosciences, Ann Arbor, MI, USA). Paired-end sequencing (150 bp) was performed on a DNBSEQ-G400 platform (MGI Tech, Shenzhen, China). After quality filtering of raw reads, clean reads were mapped against the female Nishikigoi reference genome assembly “Sh_v1.0” (NCBI accession no. BAABNH000000000) (Hosaka 2023) using Bowtie 2 (Langmead and Salzberg 2012). Unmapped reads were collected and *de novo* assembled into contigs using SPAdes (Bankevich *et al*. 2012). Contigs with a length of 5 kb or greater were selected as candidate male-specific genomic sequences and were appended to the female reference assembly “Sh_v1.0” to construct a pseudo-male reference genome. Reads from both bulk pools were re-mapped to this pseudo-male reference genome using Bowtie 2 (Langmead and Salzberg 2012). Differentially represented contigs between the female and male bulk pools based on mapping read counts were identified, yielding a final set of candidate male-specific contigs.

Similarity searches of male-specific candidate contigs were performed against the male common carp genome sequence, Ccarpio_male_YR_1.0 (GCA_060697245.1) (Chen *et al*. 2026), using the BLAST program with the default parameters (Altschul *et al*. 1997).

### Primer design and PCR screening

Based on the selected candidate contigs, 10 pairs of male-specific primers and five pairs of sex-universal (control) primers were designed (Table 1) using Primer3 (Koressaar and Remm 2007).

**Table 1.**
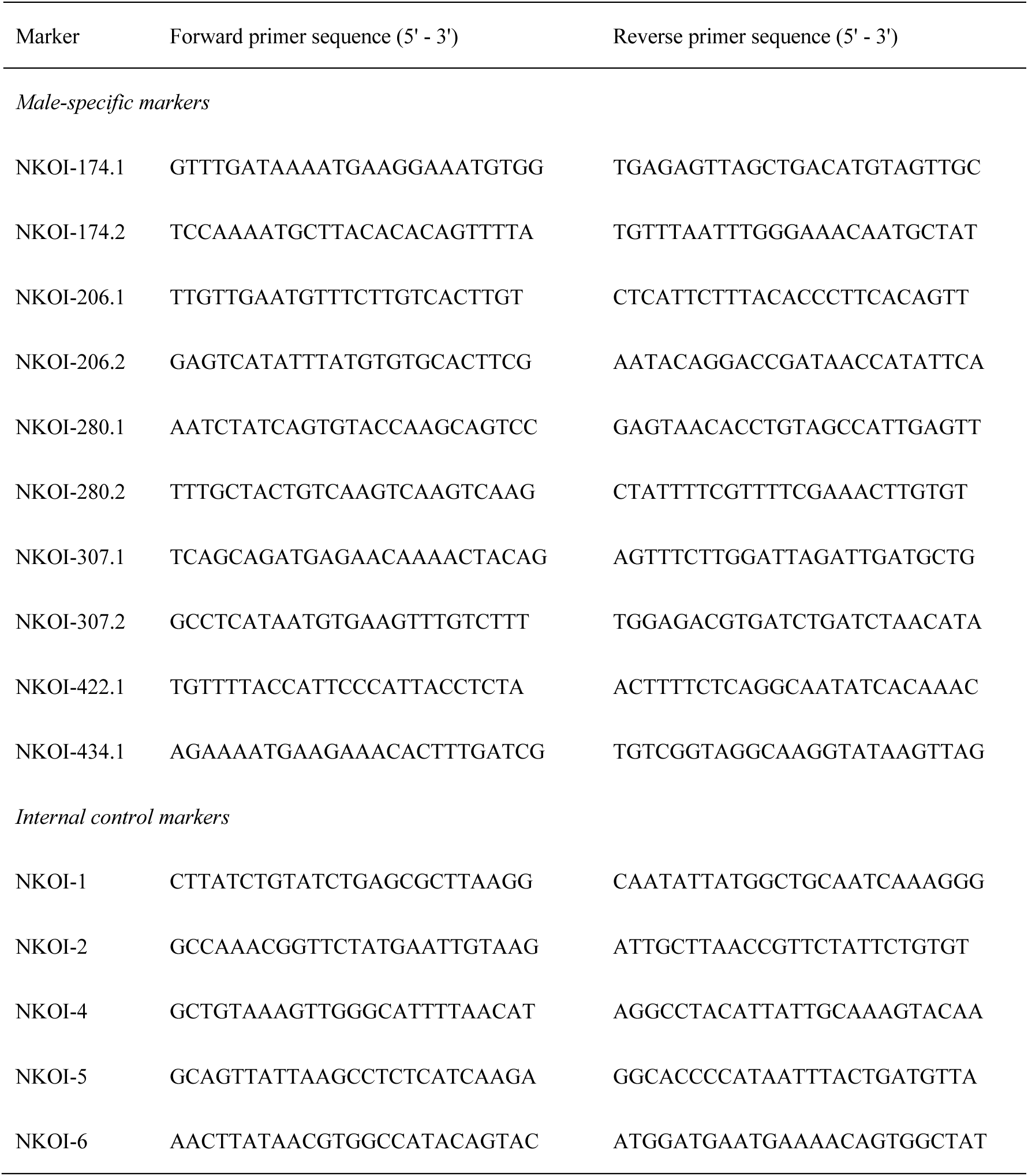
Sequences of oligonucleotides used in PCR analysis.

PCR amplification was performed using BIOTAQ DNA Polymerase (Meridian Bioscience, Cincinnati, OH, USA) with Nishikigoi genomic DNA as a template. The thermal cycling conditions were as follows: initial denaturation at 95°C for 3 min; 30 cycles of denaturation at 94°C for 30 s, annealing at 65°C for 30 s, and extension at 72°C for 1 min; followed by a final extension at 72°C for 3 min. The resulting PCR products were analyzed by 2.0% agarose gel electrophoresis and visualized with Midori Green Advance (NIPPON Genetics, Tokyo, Japan).

### Non-invasive and field-applicable sex identification

Genome DNA was extracted from epidermal mucus using GenCheck DNA Extraction Kit (Fasmac, Atsugi, Japan) and eluted in 90 µL GenCheck Buffer (Fasmac). The concentration of the extracted DNA was measured using the Qubit dsDNA HS Assay Kit (Thermo Fisher Scientific, Waltham, MA, USA) DNA was amplified with the LAMP method as described by (Takabatake *et al*. 2025). In brief, a 5-µL aliquot of the extracted DNA solution was mixed with 5 µL of the reaction mixture containing primers listed in Table 2 and heated at 63°C for 30 min to amplify DNA on a water bath. After cooling to room temperature for 5 min, 90 µL of H_2_O was added to the reaction mixture. Subsequently, 1 µL aliquot of the mixture was mixed with 15 µL of the developing buffer. A C-PAS membrane (Tohoku Bio Array), on which complementary tag sequences were printed linearly, was dipped into the solution, and target DNA was visually identified by the appearance of blue lines after 10 - 15 min.

**Table 2.**
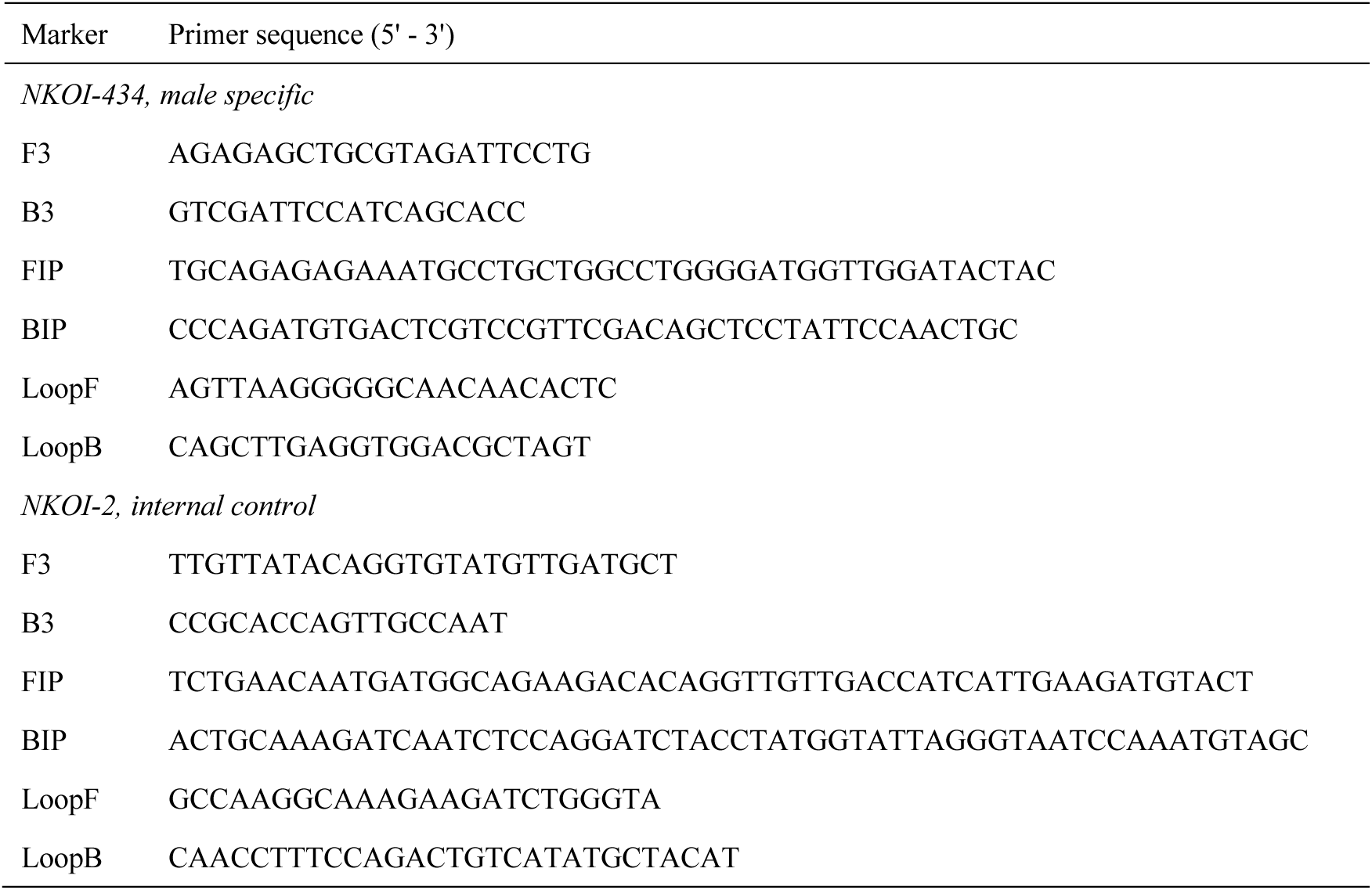
Sequences of oligonucleotides used in LAMP analysis.

## Results

### Bulked segregant sequencing analysis and selection of male-specific candidate contigs

Whole-genome shotgun sequencing of the bulked DNA samples from Kohaku generated 40.6 Gb and 30.3 Gb of high-quality clean read data for the female bulk (n = 54) and male bulk (n = 67), respectively. Mapping of these clean reads against the female reference genome assembly “Sh_v1.0” yielded alignment rates of 96.08% for the female bulk and 96.12% for the male bulk. *De novo* assembly of the unmapped reads produced 564 contigs with a length of ≥ 5 kb, spanning a total cumulative length of 3.9 Mb. These 3.9 Mb contigs were appended to the “Sh_v1.0” assembly to construct a pseudo-male reference genome, designated as “Sh_v1.0+Y”. Re-mapping of the high-quality clean reads against the “Sh_v1.0+Y” reference slightly increased the alignment rates to 96.25% for the female bulk and 96.28% for the male bulk. Differential read counts between the two bulk pools identified six contigs (NKOI-174, NKOI-206, NKOI-280, NKOI-307, NKOI-422, and NKOI-434) that showed male bulk-specific read mapping. Consequently, these six contigs were selected as candidate male-specific sequences for further marker development.

### Development and validation of male-specific PCR markers

From the six candidate male-specific contigs, 10 pairs of primers (male-specific primer candidates) were designed (Table 1). Initial PCR screening using the female and male bulk pools, as well as individual DNA samples from 11 females and 10 males randomly selected from Kohaku, demonstrated that five primer pairs (NKOI-174.1, NKOI-174.2, NKOI-307.2, NKOI-422.1, and NKOI-434.1) produced male-specific amplification products (Figure 1).

**Figure 1.**
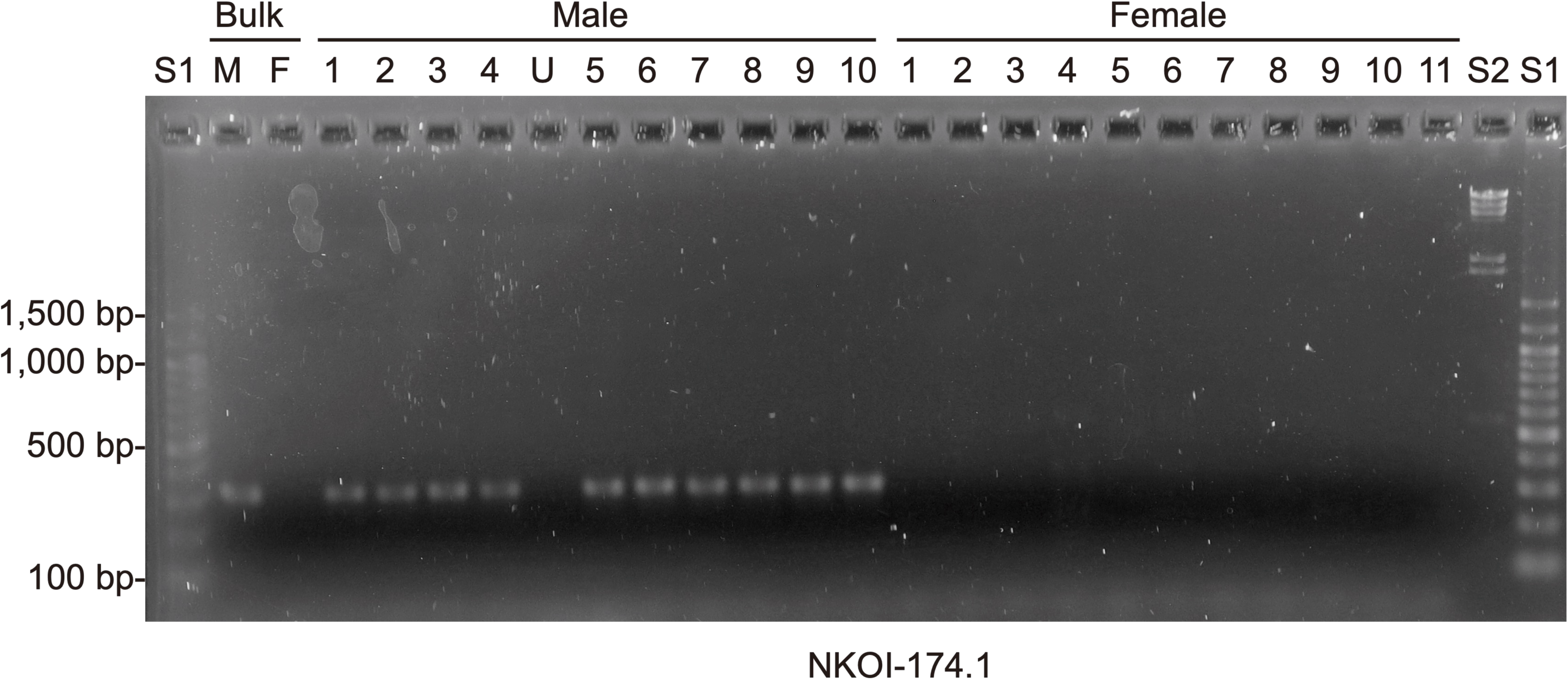
Agarose gel electrophoretic analysis of male-specific PCR amplification products. PCR products were amplified using the NKOI-174.1 primer pair from bulked male (M) and female (F) samples, as well as individual male (1–10) and female (1–11) DNA samples. “U” indicates an individual whose sex was undetermined due to immature gonads. Lanes S1 and S2 represent size markers: a 100-bp DNA ladder (Nacalai Tesque, Kyoto, Japan) and λ-HindIII digest (Life Technologies, Carlsbad, CA, USA), respectively.

These five primer pairs were further evaluated using genomic DNA from 144 Kohaku individuals. Among the 121 sex-determined individuals (54 females and 67 males), primer pair NKOI-307.2 exhibited 100% concordance between PCR amplification (presence/absence) and sex phenotypes (Table 3). Three primer pairs (NKOI-174.1, NKOI-174.2, and NKOI-422.1) each showed a discordance in a single male individual, while NKOI-434.1 failed to match the phenotypic sex in three male individuals (Table 3).

**Table 3.** Number of individuals that phenotypes matched genotypes determined by PCR male-specific markers.

| Marker | Total |  | Kohaku |  | Doitsu Karasu |  | Aka Bekko |  | Ogon |  | Magoi |  |
|---|---|---|---|---|---|---|---|---|---|---|---|---|
|  | Female | Male | Female | Male | Female | Male | Female | Male | Female | Male | Female | Male |
|  | n = 124<sup>a</sup> | n = 142 | n = 54 | n = 67 | n = 11 | n = 13 | n = 18 | n = 12 | n = 11 | n = 20 | n = 30 | n = 30 |
| NKOI-174.1 | 124 | 141 | 54 | 66 | 11 | 13 | 18 | 12 | 11 | 20 | 30 | 30 |
| NKOI-174.2 | 124 | 141 | 54 | 66 | 11 | 13 | 18 | 12 | 11 | 20 | 30 | 30 |
| NKOI-307.2 | 96 | 136 | 54 | 67 | 11 | 8 | 18 | 12 | 9 | 20 | 4 | 29 |
| NKOI-422.1 | 124 | 141 | 54 | 66 | 11 | 13 | 18 | 12 | 11 | 20 | 30 | 30 |
| NKOI-434.1 | 121 | 139 | 54 | 64 | 11 | 13 | 15 | 12 | 11 | 20 | 30 | 30 |

<sup>a</sup>Number of individuals PCR tested

To evaluate the broader applicability of the markers across different genetic backgrounds, PCR assays were subsequently conducted using the same five primer pairs in four additional populations: Doitsu Karasu (n = 32), Aka Bekko (n = 32), Ogon (n = 32), and wild common carp (Magoi, n = 60). For the sex-determined individuals in these groups, comprising Doitsu Karasu (11 females, 13 males), Aka Bekko (18 females, 12 males), Ogon (11 females, 20 males), and Magoi (30 females, 30 males), three primer pairs (NKOI-174.1, NKOI-174.2, and NKOI-422.1) demonstrated 100% concordance between amplification and phenotypic sex (Table 3). However, NKOI-174.2 displayed variation in PCR amplicon size among certain samples. In contrast, NKOI-307.2 achieved complete concordance in Aka Bekko but exhibited discordant amplification in five Doitsu Karasu males, two Ogon females, and 26 females and one male of Magoi (Table 3). Primer pair NKOI-434.1 matched phenotypic sex in Doitsu Karasu, Ogon, and Magoi, but showed discordance in three Aka Bekko female individuals (Table 3).

Based on these comprehensive validation results among the 266 individuals for which phenotypic sex was determined (Table 3), two primer pairs (NKOI-174.1 and NKOI-422.1) which achieved an exceptional accuracy rate of 99.6% (265/266 individuals matched), were selected as the primary male-specific diagnostic markers for Nishikigoi. NKOI-174.2 also exhibited 99.6% concordance (265/266 individuals matched) in total but not selected due to the variation in PCR amplicon size. Additionally, NKOI-434.1 proved highly reliable as a secondary marker, demonstrating a strong accuracy rate of 97.7% (260/266 matched).

### Development and screening of internal control PCR markers

To establish positive controls for PCR assays, five candidate sex-universal primer pairs designed from five contigs (NKOI-1, 2, 4, 5, and 6) were evaluated (Table 4). Initial screening across 96 individuals (Doitsu Karasu, Aka Bekko, and Ogon; n = 32 each) identified NKOI-2 with 100% amplification success (Table 4). In contrast, NKOI-1, 4, 5, and 6 failed in 4, 4, 15, and 19 individuals, respectively, during initial screening.

**Table 4.** Number of individuals that DNA were amplified by PCR internal control markers.

| Marker | Total | Doitsu Karasu | Aka Bekko | Ogon | Kohaku | Magoi |
|  | Amplified / Tested<sup>a</sup> | n = 32<sup>b</sup> | n = 32 | n = 32 | n = 144 | n = 60 |
|---|---|---|---|---|---|---|
| NKOI-1 | 92 / 96 | 28 | 32 | 32 | n.a. | n.a. |
| NKOI-2 | 299 / 300 | 32 | 32 | 32 | 144 | 59 |
| NKOI-4 | 92 / 96 | 29 | 31 | 32 | n.a. | n.a. |
| NKOI-5 | 81 / 96 | 32 | 29 | 20 | n.a. | n.a. |
| NKOI-6 | 77 / 96 | 31 | 14 | 32 | n.a. | n.a. |

<sup>a</sup>Number of individuals that DNA were amplified by PCR out of number of individuals PCR tested
<sup>b</sup>Number of individuals PCR tested

Further validation of the top three candidates across 144 Kohaku and 60 Magoi (n = 204) yielded 99.5% (203/204) for NKOI-2, failing in only a single Magoi specimen (Table 4). Overall, NKOI-2 was selected as a reliable internal control.

### DNA diagnostics for Nishikigoi sex identification

For the DNA diagnostic assay, the NKOI-434 and NKOI-2 sequences were selected because functional LAMP primers could be successfully designed targeting these two loci (Table 2). We evaluated the performance of this diagnostic method using genomic DNA from 32 individuals (15 males, 15 females, and two sex-undetermined individuals) whose NKOI-434.1 genotypes had been previously determined by standard PCR (yielding 16 male-type and 16 female-type results). The genomic DNA was amplified by LAMP, followed by a 1:10 dilution with H_2_O and mixing with the developing buffer. Subsequently, amplicons were detected using C-PAS. As expected, positive signals for both the male-specific sequence and the internal control were detected in all 15 male samples, as well as in one PCR-positive sex-undetermined individual (Figure 2A). In contrast, only the internal control signal was observed in 13 female samples and one PCR-negative sex-undetermined individual, whereas positive signals for both target sequences were detected in the remaining two PCR-negative female samples (i.e., false positives).

**Figure 2.**
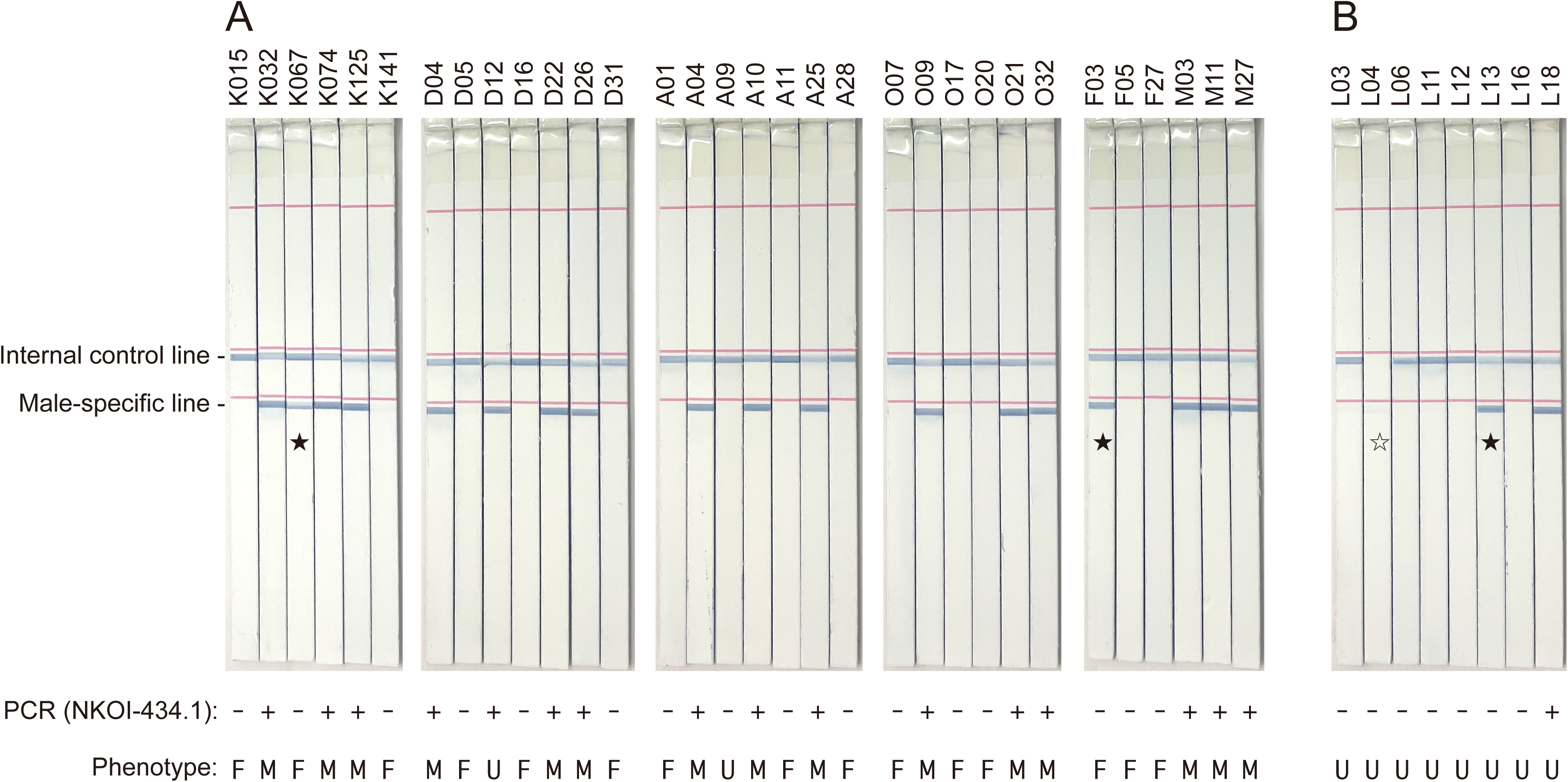
DNA diagnostics for Nishikigoi sex identification using chromatography-printed array strip (C-PAS). Internal control and male-specific signals are detected at each line position. (A) DNA samples from four Nishikigoi varieties (K: Kohaku; D: Doitsu Karasu; A: Aka Bekko; and O: Ogon) and wild common carp (F and M: Magoi). (B) DNA extracted non-invasively from the epidermal mucus of living fish samples (L). PCR results using the NKOI-434.1 primer pair (+: positive; and -: negative) and sex phenotypes (M: male; and F: female) are shown at the bottom. Solid and open stars indicate false-positive and failed samples, respectively.

Furthermore, we applied this diagnostic platform to eight living individuals whose sex phenotypes were unknown. The concentration of genomic DNA extracted non-invasively from the epidermal mucus ranged from 0.4 to 5.5 ng/µL, with an average of 1.7 ng/µL. Following LAMP amplification, 1:10 dilution, and C-PAS detection, positive signals for both the male-specific sequence and internal control were detected in two individuals (one PCR-positive and one PCR-negative; Figure 2B). Only the internal control signal was observed in five PCR-negative individuals, whereas no signal was detected for either line in the remaining PCR-negative sample, likely due to insufficient template DNA quantity or amplification failure.

### Genome positions of the male-specific candidate contigs

During the preparation of this manuscript, a chromosome-scale genome assembly of a male common carp was published (Chen *et al*. 2026), revealing that a male-specific gene encoding *Pangu*, a helix-loop-helix (HLH) domain peptide, is located at 21.4 Mb on chromosome 25B. Therefore, we investigated the genomic positions of the six male-specific candidate contigs identified in this study. All six contigs exhibited high sequence similarity to a 250-kb region (21.25–21.50 Mb) on chromosome 25B, corresponding to the genomic segment containing *Pangu* (Figure 3). Interestingly, the NKOI-174 contig spanned the entire Y-specific fragment and harbored the complete coding sequence for the 76-amino-acid *Pangu* gene. The physical distances between the two markers and the *Pangu* gene were 0.5 kb for NKOI-174.1 and 3.1 kb for NKOI-174.2.

**Figure 3.**
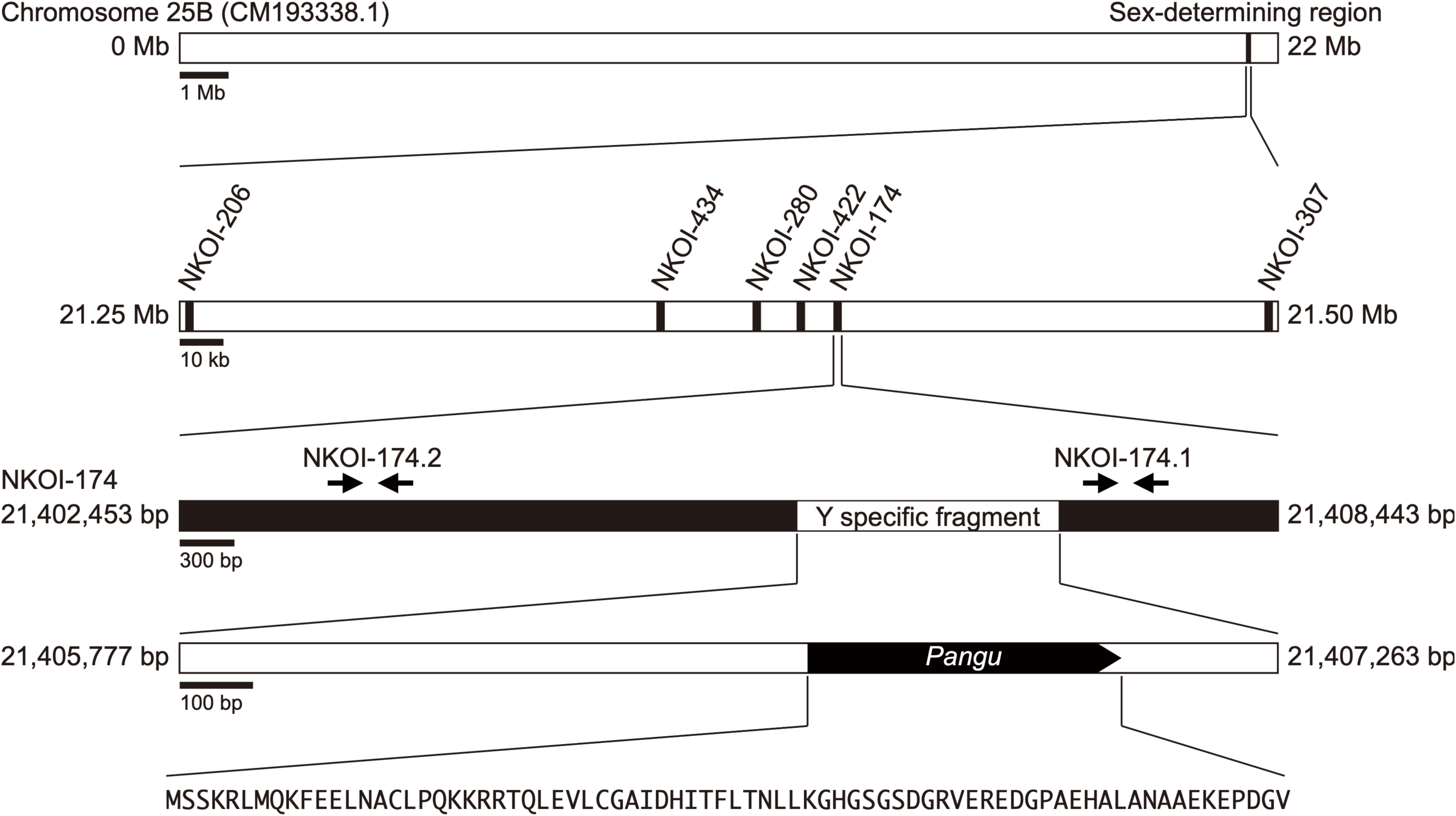
Physical map of the male-specific region on carp chromosome 25B. The six candidate contigs (NKOI-206 to NKOI-307) are located within a 250-kb region around the 21.25–21.50 Mb locus on chromosome 25B. Among them, the NKOI-174 contig contains two PCR marker sequences (NKOI-174.1 and NKOI-174.2) and the complete coding sequence of the male-specific gene (*Pangu*) within the Y-specific region. The predicted amino acid sequence of *Pangu* is shown at the bottom. The locations of the “Sex-determining region”, “Y specific fragment”, and *Pangu* were mapped according to Chen et al. (2026).

## Discussion

The BSA combined with whole-genome shotgun sequencing successfully identified novel candidate male-specific sequences, enabling the development of highly accurate PCR-based sex-identification markers in Nishikigoi (Table 3, Figure 1). In particular, two primary primer pairs, NKOI-174.1 and NKOI-422.1, exhibited near-perfect diagnostic accuracy, with amplification results completely matching phenotypic sex across all evaluated individuals, except for a single male Kohaku specimen (99.6% concordance; Table 3). These results demonstrate that these markers provide a highly reliable tool for molecular sex identification in Nishikigoi. The strong association of these markers with phenotypic sex suggests that they are likely linked to the sex-determining region. Indeed, the six male-specific contigs were located within the 250-kb genome region on the chromosome 25B including *Pangu* (Chen *et al*. 2026) (Figure 3), suggesting that the sex determination in common carp and Nishikigoi was governed by the same genetic mechanism.

Because all five male-specific PCR markers function as dominant markers producing positive amplicons exclusively in males, they possess inherent technical limitations. Dominant markers cannot distinguish homozygous from heterozygous genotypes. More critically, any PCR failure caused by insufficient DNA quantity, PCR inhibition, or reagent degradation could result in false-negative results, wherein a true male individual is incorrectly diagnosed as female. To overcome this limitation and ensure diagnostic accuracy, we developed and validated sex-universal PCR markers targeting the nuclear genome (Table 4). Although universal primer sets targeting mitochondrial DNA, such as MiFish and COI (Ivanova *et al*. 2007, Miya *et al*. 2015), are also available, we chose not to use them because their exceptionally high copy number relative to nuclear DNA could potentially increase the risk of false-positive amplifications.

In our diagnostic evaluation, a total of 40 samples were tested (Figure 2). Signal amplification was successfully achieved in 39 samples, among which 36 samples (92.3%) exhibited identical results between the PCR and C-PAS analyses. Discrepancies were observed in three samples, all of which were identified as female by PCR but yielded male-pattern results with C-PAS (i.e., discordance due to false positives in C-PAS). The remaining single mucus sample produced no detectable signals, likely due to an insufficient amount of template DNA. In field settings, the C-PAS method is highly advantageous over standard PCR because it does not require specialized molecular biology equipment. The exact reasons for these false positives remain to be fully elucidated, but several plausible explanations exist. First, intrinsic differences in sensitivity and specificity between the LAMP and PCR assays, including primer specificity, could contribute to the observed discrepancy (Foo *et al*. 2020, Khan *et al*. 2018). Second, because male and female fish were co-housed in the same holding tanks, trace amounts of male DNA may have contaminated the female samples as environmental DNA (eDNA) (Ficetola *et al*. 2008). Given the exceptionally high sensitivity of isothermal LAMP amplification, such trace amounts of template DNA might be sufficient to generate a signal in LAMP, whereas they would remain below the detection threshold for standard PCR. Nevertheless, from a practical standpoint in commercial breeding, misidentifying a female as a male (false positive) is generally far more acceptable than misidentifying a male as a female (false negative), as it merely leads to the conservative exclusion of an individual rather than an unwanted contamination of sex-selected broodstock.

While this study successfully established practical sex-identification markers, the male-specific genomic sequences identified here also provide valuable genomic resources for further characterizing the Y-chromosome sex-determination system of *C. carpio*. The markers and sequences identified in this study therefore provide a foundation for practical, non-invasive sex identification in Nishikigoi and for further investigation of the genomic basis of sex determination in common carp.

## Data availability

Sequence data generated in this study are available from the DDBJ BioProject database under the accession number PRJDB46060.

## Acknowledgements

We are grateful to Mr. T. Komatsu (PacBio Japan) for facilitating this work. We also thank Y. Gomi, C. Minami, H. Tsuruoka, Y. Kishida, A. Obara, and A. Watanabe (Kazusa DNA Research Institute) for technical assistance. This study was supported by the Kazusa DNA Research Institute Foundation. Gemini (Google) was used to assist with English language editing and translation during the preparation of this manuscript. The authors reviewed and took full responsibility for the final content of the manuscript.

